# Neutrophils drive sexual dimorphism in experimental periodontitis

**DOI:** 10.1101/2024.11.27.625678

**Authors:** Kelsey Martin, Maxwell Mianecki, Victoria Maglaras, Asfandyar Sheikh, Muhammad H. A. Saleh, Ann M. Decker, Joseph T. Decker

## Abstract

The motivating premise of this study is to improve the treatment of periodontal disease by elucidating sex-specific mechanisms of periodontal disease progression. Men and women experience inflammation in fundamentally different ways and understanding the sex-specific biology leading to inflammation and bone loss in the periodontium will inevitably improve patient outcomes. We therefore examined clinical and immunological differences in the progression of periodontal disease using the ligature-induced periodontitis model. Periodontitis was induced in male and female C57BL/6j mice by tying a 5-0 silk suture around the left maxillary second molar. The ligature was left on for 7 or 21 days at which point maxillae were characterized for bone loss by μCT or immune infiltrate by flow cytometry. Neutrophil depletion was accomplished through systemic administration of a Ly6G antibody. Conditions were compared using two-way ANOVA with Tukey’s multiple comparison correction from n≥5 animals. Ligature-induced periodontitis led to alveolar bone loss at both 7 and 21 days in both female and male mice. Males and females had approximately the same amount of linear bone loss 7 days post-ligature placement, while male mice had significantly more linear bone loss by day 21. Male mice had significantly more immune cells in their maxillae 7 days post ligature placement compared to female mice. Both male and female mice showed a shift in immune populations towards neutrophils, with no significant difference between males and females. Neutrophil counts were significantly elevated in male mice on day 7 but not day 21, while female mice did not have any statistically significant changes in neutrophil counts. Neutrophil depletion using a Ly6G antibody limited bone loss in male but not female mice relative to isotype antibody-treated controls. Analysis of single-cell sequencing data from human patients with periodontitis showed differences in neutrophil phenotypes that were also observed in a mouse model of periodontitis. Together, these data suggest a mechanistic role for neutrophil inflammation in sexual dimorphism in periodontitis.

## Introduction

Periodontal disease is a debilitating inflammatory condition within the supporting tissues of the teeth (including gingiva, alveolar bone, and the periodontal ligament) that leads to significant morbidity, systemic consequences, and economic difficulties for patients^1–7^. Periodontal disease is extremely common; a recent NHANES stratified multistage probability sample of the civilian noninstitutionalized population assessment reported that 42% of the US population has some type of periodontitis (severe, moderate, or mild), including 7.8% being severe periodontitis as defined by the CDC/AAP periodontitis case definitions^8^. Furthermore, the disease burden of periodontal disease has significant consequences for overall health, as periodontal disease has been associated with numerous comorbidities, including diabetes, cardiovascular disease, adverse pregnancy outcomes, rheumatoid arthritis, and chronic obstructive pulmonary disease^4–6^. Clearly, understanding the patient-specific mechanistic underpinnings of periodontal disease are essential for both oral and systemic health.

Biological sex is one such patient-specific mechanism that may contribute to the overall prevalence and severity of periodontal disease. Periodontal disease is more common in men compared to women^8–13^. Additional studies mechanistically attribute this sex-biased presentation to behavior patterns, habits, attitudes, which form a part of environmental risk factors rather than constitutional factors^14, 15^. However, there is newer evidence that this previously held belief is incorrect and there exists a biological foundation to the sexual dimorphism epidemiological observation^16–18^, and social determinants can only explain some of the observed sexual dimorphism in the prevalence and severity of periodontal disease^19^.

Sexual dimorphism has been widely observed in the clinical prevalence of periodontitis, yet very few mechanistic studies have been performed to elucidate the underlying mechanisms for these differences. Therefore, a need exists to dissect the specific biology underpinning sexual dimorphism in periodontitis. The primary goal of this study was, therefore, to define the contribution of biological sex to the progression of periodontal disease in a preclinical model system and identify potential immunological mechanisms underpinning sexual dimorphism in this disease.

## Methods & Materials

### Animals

This study used male and female C57Bl/6J mice or Padi4^-\-^ mice on a C57Bl/6J background (Jackson Laboratories, Bar Harbor, Maine). Mice were maintained according to institutional animal care and use guidelines, and experimental protocols were approved by the Institutional Animal Care and Use Committee of the University of Michigan. Mice were housed in groups of 2- 5 under specific pathogen-free conditions.

### Ligature-induced Periodontitis

Periodontitis was induced in mice using the previously described ligature-induced periodontitis model^20^. Mice six to eight weeks in age were anesthetized using a ketamine/xylazine cocktail following isoflurane vaporizer induction, and a 5-0 silk suture (Roboz, Gaithersburg, MD) was tied around the second maxillary molar. The ligature was left on for 7 or 21 days before being euthanized to evaluate periodontitis progression. Mice were excluded if they experienced significant trauma during ligature placement, were found to be missing a ligature at the time of euthanasia, or lost a tooth to periodontal disease. All experiments were conducted with an n≥5.

### Micro-computed Tomography

Maxillae were harvested at euthanasia, fixed in 10% buffered formalin for 24-72 hours at 4 °C on a rocker, and stored in phosphate-buffered saline. Specimens were scanned using a micro-CT system (µCT100 Scanco Medical, Bassersdorf, Switzerland). Scan settings for maxillae were: 34 mm sample holder, voxel size 18 µm, 70 kVp, 114 µA, 0.5 mm AL filter, and integration time 500 ms. Images were analyzed by a single calibrated blinded examiner using Dragonfly (Comet Technologies). Distance from the cemento-enamel junction to the alveolar ridge was measured at four sites (mesial, buccal, distal, and palatal) at the midline of the second molar^21^. Bone loss was calculated as the difference between average bone levels between ligated and unligated sides of the same mouse. Volumetric measurements were calculated as the volume from 10 slices below the cemento-enamel junction (approximate height of healthy bone) to the alveolar bone crest^21^.

### Histology

Maxillae and tibia were fixed in 10% buffered formalin for 24-72 hours at 4 °C on a rocker and stored in 70% ethanol. Samples were placed into labeled tissue cassettes and decalcified in a 14% EDTA solution for 2 weeks. All cassettes were processed in a short amount of time using a VIP tissue processor. Samples were paraffin-embedded using a Leica paraffin embedding machine, and sagittal sections were cut at 5 µm. Sections were stained with hematoxylin and eosin for bone and cellular identification.

### Flow Cytometry

Maxillae were harvested and kept in RPMI media until all samples were collected. Maxillae were cut in half through the palate, at the end of the hard palate, and before the incisor. Maxillae blocks were transferred into digestion media containing DNase1 (0.1 mg/ml; Roche) and Liberase (0.1 mg/ml; Sigma-Aldrich) and incubated at 37 °C for 30 minutes, minced, incubated for another 30 minutes, and digestion stopped with 2700 µL complete media. Tissue and media were filtered using a 70 µm cell strainer, centrifuged for 5 minutes at 4 °C, the supernatant was decanted, and cells resuspended before being transferred to micro-centrifuge tubes. Peripheral blood (neutrophil depletion) was lysed using 2.5 mL of ACK lysis buffer (Life Technologies Corporation) and incubated for 10 minutes before centrifugation and washing. Samples were incubated for 10 minutes with flow cytometry antibodies. The following antibodies and dilutions were used:

PE/Cy7- CD49b 1:250, AlexaFluor 700- F4/80, PerCP/Cy5.5- CD4 1:250, PerCP- CD44 1:250, BV711- CXCR4 1:250, BV510- CD19 1:250, BV605- NK1.1 1:250, FITC- CD11b 1:250, BV785- CD11c 1:250, PE- CD45R/B220 1:250, BV570- CD62L, BV421- Ly6G 1:250; Biolegend; BV480-CD8a 1:250, NovaFluor Yellow 730- CD3a 1:50, Invitrogen; APC- Ly6C 1:250; BD Bioscience, and live/dead stain (Fixability Dye eFluor450 1:1000; eBioscience). Samples were analyzed using a Cytek Northern Lights (3 laser; V-B-R). Data was unmixed using single color controls using SpectroFlow software. Data was processed using FlowJo v 10.8.1 (Becton Dickinson). The gating scheme used to identify cell types can be seen in **FIG S1**.

### Neutrophil Depletion

200µL of diluted anti-Ly6G (Bio X Cell, clone 1A8) or Rat 1gG2a isotope control (0.168 µL/mL) was injected intraperitoneally every other day for 22 days, beginning on Day -1. Retro-orbital blood was collected on Day 4 to confirm neutrophil depletion via flow cytometry.

### Single-cell sequencing analysis

Publicly available, human single cell RNA sequencing datasets – GSE164241^22^ and GSE171213^23^– of patients with and without periodontitis were collected for analysis. Patient files included for evaluation were those collected from periodontal tissues of adults (≥ 18 years of age) in good general health and not within 3 months of taking anti-inflammatory or anti-bacterial drugs. Inclusion criteria were similar between the two studies, with patients in the moderate to severe periodontitis groups having probing depths over 5 mm and bleeding upon periodontal probing. Analyses were performed in R, utilizing the Seurat v3 package^24^. Dataset files were first downloaded from GEO Accession Viewer. GSE164241 files were input using ‘Read10X’ and GSE 171213 files were input utilizing ‘read_table’ followed by ‘read.delim’ to create a Seurat-compatible format using the ‘CreateSeuratObject’ function. Files were merged, using the ‘merge’ function into groups based on biological sex and periodontal condition – healthy female, healthy male, periodontitis female, and periodontitis male – for subsequent evaluation. Quality control, normalization, and integration were performed as previously described^22, 24^. Visualization and clustering were accomplished using ‘ScaleData’ to account for cell cycle followed by ‘FindNeighbors’ and ‘FindClusters’. Markers for each cluster, identifying unique cell types, were identified using ‘FindMarkers’, and in totality using ‘FindAllMarkers’ (only.pos = TRUE, logfc.threshold = 0.25, min.pct = 0.5), based on differential gene expression. UMAP and heatmap plots were produced to reflect highly expressed markers of interest by cluster. Relative expression plots of cell-specific subtypes were generated to discern expression differences between the four groups. Cell-cell communication networks were inferred using CellChat^25^. Gene ontology was performed using MetaScape^26^.

### Statistical Analysis

Statistical analysis was performed using GraphPad Prism version 10.3.0 for Windows (GraphPad Software, Boston, MA). Bone loss measurements and flow cytometry results were compared using two-way ANOVA followed by Tukey’s multiple comparison correction unless otherwise noted. Single comparisons were analyzed by *t*-test. A p-value<0.05 was used for statistical significance in all studies. All experiments were performed on n≥5 mice.

## Results

### Male and female mice have different amounts of bone loss in chronic periodontitis

We began our study of sexual dimorphism in experimental periodontitis by comparing the progression of ligature-induced periodontitis (LIP) in male and female C57Bl/6J mice (**FIG 1**). Male and female mice had their teeth ligated and alveolar bone levels at the site of the ligature compared at days 7 and 21 post ligature placement (**FIG 1A**). These time points were selected to mimic the progression of periodontal disease in humans^27^, with the day 7 time point approximating acute inflammation and the day 21 approximating chronic inflammation. Male and female mice had similar levels of alveolar bone prior to ligature placement as measured by µCT (**FIG 1B, FIG 1C**). Ligature-induced periodontitis led to alveolar bone loss at both 7 and 21 days in both female and male mice (**FIG 1D**). Males and females had approximately the same amount of linear bone loss in the 7-day model of acute periodontitis (0.20 mm male, 0.22 mm female, p=0.79 **FIG 1E**). Male mice had significantly more linear bone loss in the 21-day model of chronic periodontitis (0.60 mm male, 0.46 mm female, p<0.0001, **FIG 1E**). Measuring the volume of lost bone showed similar results, with similar bone loss 7 days post ligature placement (0.33 mm^3^ male, 0.33 mm^3^ female, p>0.9999) and greater bone loss in male mice 21 days post ligature placement (0.90 mm^3^ male, 0.67 mm^3^ female, p<0.001, **FIG 1F**).

**FIG 1:**
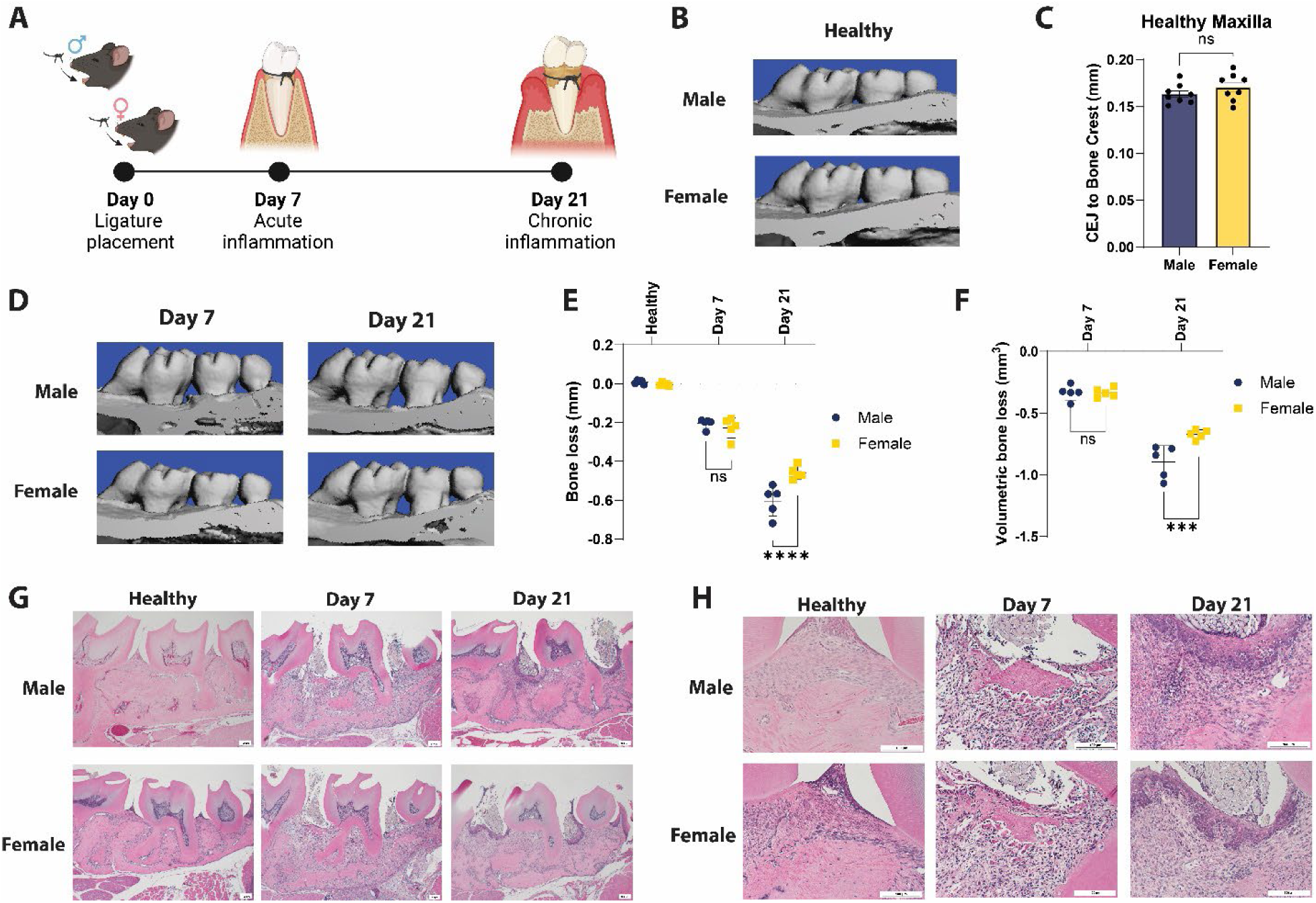
Sexual dimorphism in the clinical phenotype of experimental periodontitis. A) Time course of ligature induced periodontitis. B) MicroCT images of healthy maxillae in male and female mice. C) Cemento-enamel junction (CEJ) to alveolar bone crest distances in healthy male and female mice. D) MicroCT images of day 7 and day 21 maxillae in male and female mice. E) Linear bone loss measurements in male and female mice at the healthy, day 7, and day 21 time points. F) Volumetric bone loss in male and female mice at the day 7 and day 21 time points. G) Histology images of entire ligature site in male and female mice. Scale bar = 100µm H) High magnification image of mucosal surfaces between the first and second molar. Scale bar = 50 µm. ns= not significant. ***p<0.001, ****p<0.0001, two-way ANOVA with Tukey multiple comparison correction. Data are presented with bars indicating the mean value ± standard deviation.

In male and female healthy mice there is a shallow gingival sulcus around the teeth (**FIG 1G, FIG 1H**). Histologically the gingival sulcus is lined by the sulcular epithelium, attached at the most apical aspect to the enamel surface. The alveolar bone is well-defined with distinct and intact borders. At day 7 time points histological staining depicts the junctional epithelium has detached from the tooth surface, forming an intra-epithelial cleft with increased permeability. The pocket epithelium demonstrates a high infiltration, particularly of the epithelial ridges, with neutrophilic granulocytes and lymphocytes. The alveolar bone is not clearly linear on the coronal and lateral borders. At day 21 time points, proliferation of the epithelial ridges into aspects of the soft connective tissue is noted with very thin regions between these ridges. Focal micro-ulcerations of the epithelial ridges are noted at the free surface of the pocket epithelium. Additional interdental alveolar bone dimensional loss has occurred beyond 50% of the length of the root surface.

### Neutrophil infiltrate is exaggerated in male mice with periodontitis

The inflammatory response is one of the primary drivers of bone loss in periodontitis. We therefore characterized the changes in the immune cell populations during LIP male and female mice, with the hypothesis that male and female mice would have differential recruitment of immune cells (**FIG 2**). Male and female mice had similar numbers of immune cells in the maxilla prior to ligature placement (**FIG 2A**). Male mice had significantly more immune cells in their maxilla 7 days post ligature placement relative to female mice (p<0.05); this difference in total immune cell count was resolved by day 21. Both male and female mice showed a shift in immune populations towards Cd11b^+^ Ly6G^+^ neutrophils, with no significant difference between males and females (**FIG 2B**). Delineating the number of different populations of immune cells showed significantly more CD11b^+^ Ly6G^+^ neutrophils (p<0.01) and CD19^+^ B cells (p<0.05) in male maxillae 7 days post ligature placement (**FIG 2C**). The total number of neutrophils, but not B cells, was elevated in male maxillae 21 days post ligature placement compared with female mice (p<0.0001, **FIG 2D).** We further analyzed changes in neutrophil infiltrate over time (**FIG 2E, FIG 2F, FIG 2G**). Male mice showed a significant trend towards increased neutrophil fraction of immune cells (fraction of CD45+ cells) over time, while female mice had a similar, yet not statistically significant, trend (**FIG F**). Both male and female mice showed a peak in neutrophil counts in the maxilla at day 7, with only male mice having a statistically significant change from healthy (p<0.001).

**FIG 2:**
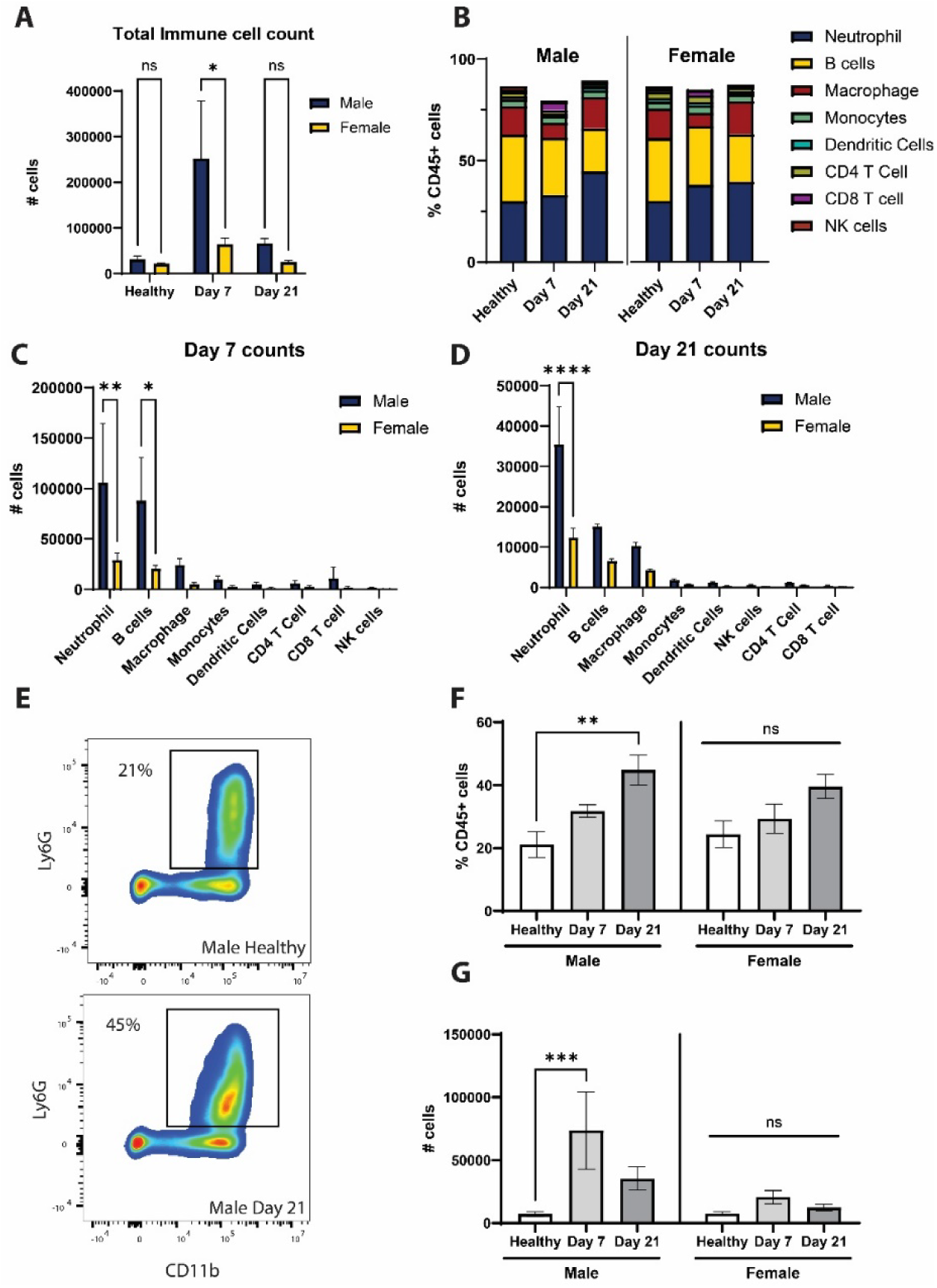
Increased neutrophil infiltrate in male mice with periodontitis. A) Total immune cell counts in the maxilla in male and female mice at different time points of LIP. B) Proportions of CD45^+^ cells by immune cell type in the maxilla. C) Counts of different immune cell populations in the maxilla at day 7. D) Counts of different immune cell populations in the maxilla at day 21. E) Example flow plot showing increased neutrophil infiltrate in the day 21 male maxilla. F) Trends towards increasing neutrophil fraction over time in the maxilla of male but not female mice. G) Peak neutrophil infiltrate at day 7. NS = not significant. * p<0.05, **p<0.01, ***p<0.001, ****p<0.0001, two-way ANOVA with Tukey’s multiple comparison correction

**FIG 3:**
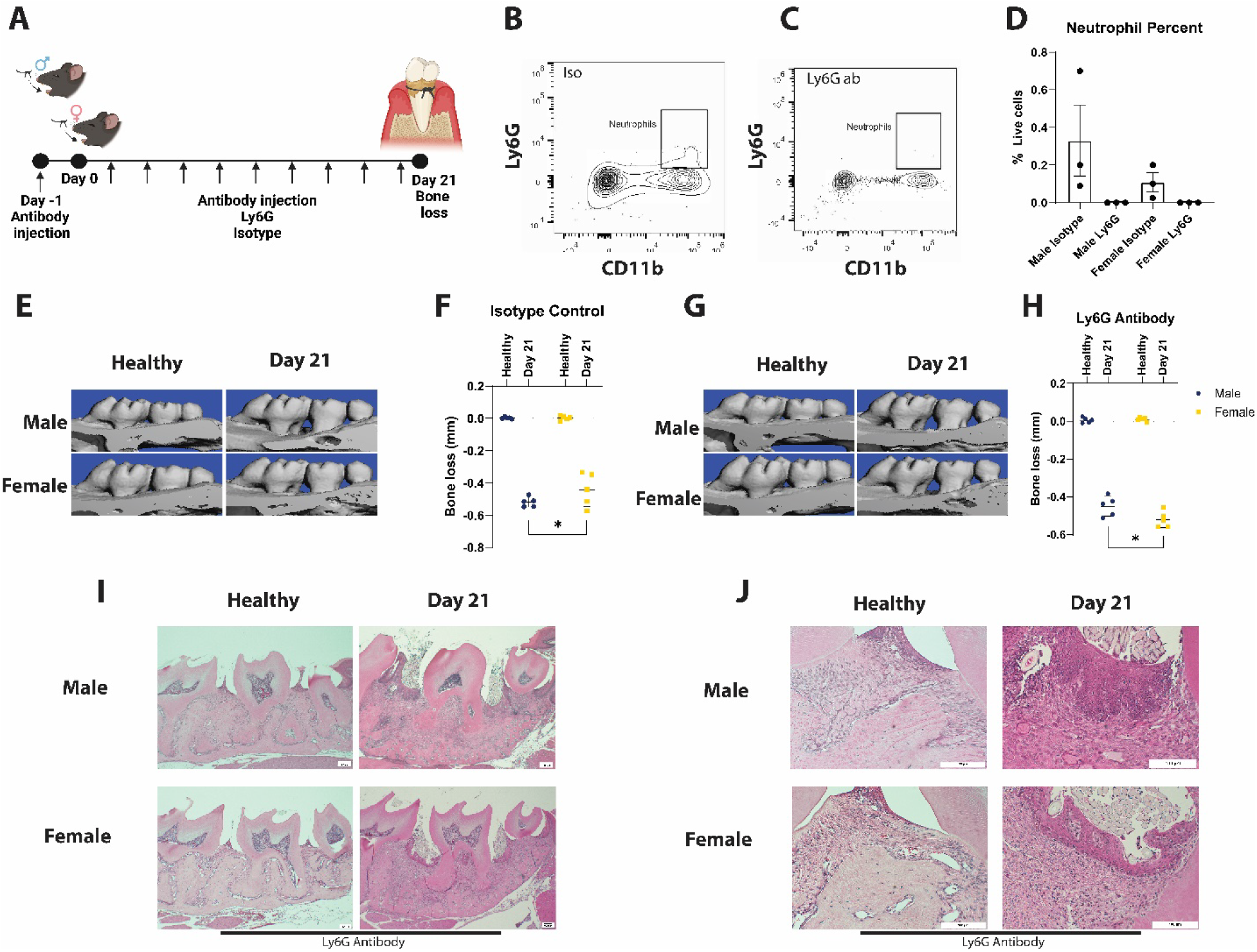
Neutrophil depletion differentially affects male and female mice. A) Experimental plan for depletion of neutrophils during LIP. B, C) Flow plots showing depletion of neutrophils in the blood of Ly6G antibody treated mice (C) but not isotype control treated mice (B). D) Quantification of neutrophil fraction in the blood during antibody treatment. E) MicroCT images showing bone loss in isotype control mice at day 21. F) Quantification of bone loss showing sexual dimorphism in isotype control treated mice with LIP. G) MicroCT images showing bone loss in Ly6G antibody treated mice. H) Quantification showing differential bone loss between male and female mice with LIP treated with Ly6G antibody. I) Low magnification histology image showing ligature site or Ly6G antibody treated mice. Scale bar = 100µM. J) High magnification image of the ligature site in Ly6G antibody treated mice. Scale bar = 50 µM. *p<0.05, two-way ANOVA.

### Neutrophil depletion differentially affects male and female mice with periodontitis

Since neutrophil inflammation persists in male mice relative to female mice, we next examined the sexual dimorphism in bone loss during periodontitis in mice depleted of neutrophils (**FIG 3A**). Systemic treatment with a Ly6G antibody successfully depleted neutrophils in the blood of both male and female mice relative to isotype antibody-treated controls (**FIG 3B, FIG 3C, FIG 3D**). Mice treated with an isotype antibody showed similar amounts of bone loss relative to untreated controls, with significantly less bone loss in female mice compared with male mice at the day 21 time point (0.51 mm male, 0.44 mm female, p<0.05, **FIG 3E**, **FIG 3F**). Sexual dimorphism was also observed in mice treated with Ly6G antibody, with female mice losing more bone than male mice (0.45 mm male, 0.52 mm female, p<0.05, **FIG 3G**, **FIG 3H**). Histological examination of the ligature sites of antibody depleted mice showed similar levels of tissue inflammation between male and female mice, including infiltrate of inflammatory immune cells (**FIG 3I)** and inflammation in the mucosal tissue abutting the ligature (**FIG 3J**).

### Human periodontitis patients have sexually dimorphic neutrophil signaling

Depletion of neutrophils differentially affected male and female mice. We therefore hypothesized that sexual dimorphism in neutrophil phenotype existed in periodontitis. We therefore examined sex differences in neutrophil phenotype in human periodontitis (**FIG 4**). To accomplish this we aggregated publicly available single-cell RNA sequencing data (scRNAseq data) of individuals with and without periodontitis. Clustering this data using the Seurat algorithm^24^ found 22 unique cell types within this data set (**FIG 4A**). Granulocytes were identified as a cluster of cells with high expression of *CSF3R*, *FCG3RB, IL1B,* and *C5AR1* (**FIG 4B**). Next, we conducted cell-cell communication analysis using CellChat^25^. Stromal elements in the oral mucosa (endothelial cells, fibroblasts, keratinocytes) were the primary sources of signaling molecules in both male and female periodontitis patients (**FIG 4C**). We next examined differences in signaling strength between paths in male and female patients (**FIG 4D**). The SELL, CXCL, and COLLAGEN pathways were the most differential pathways based on inferred signaling strength. Plotting significantly contributing receptor-ligand interactions showed differences in incoming signals through *CD44* and *CXCL* receptors *CXCR1*, *CXCR2*, and *CXCR4* (**FIG 4E**). Outgoing differences were observed *SELL*, *CXCL1*, and *CXCL8* (**FIG 4F**). CXCL1-ACKR1 interactions were specific to female neutrophils (outgoing from neutrophils to endothelial cells).

**FIG 4:**
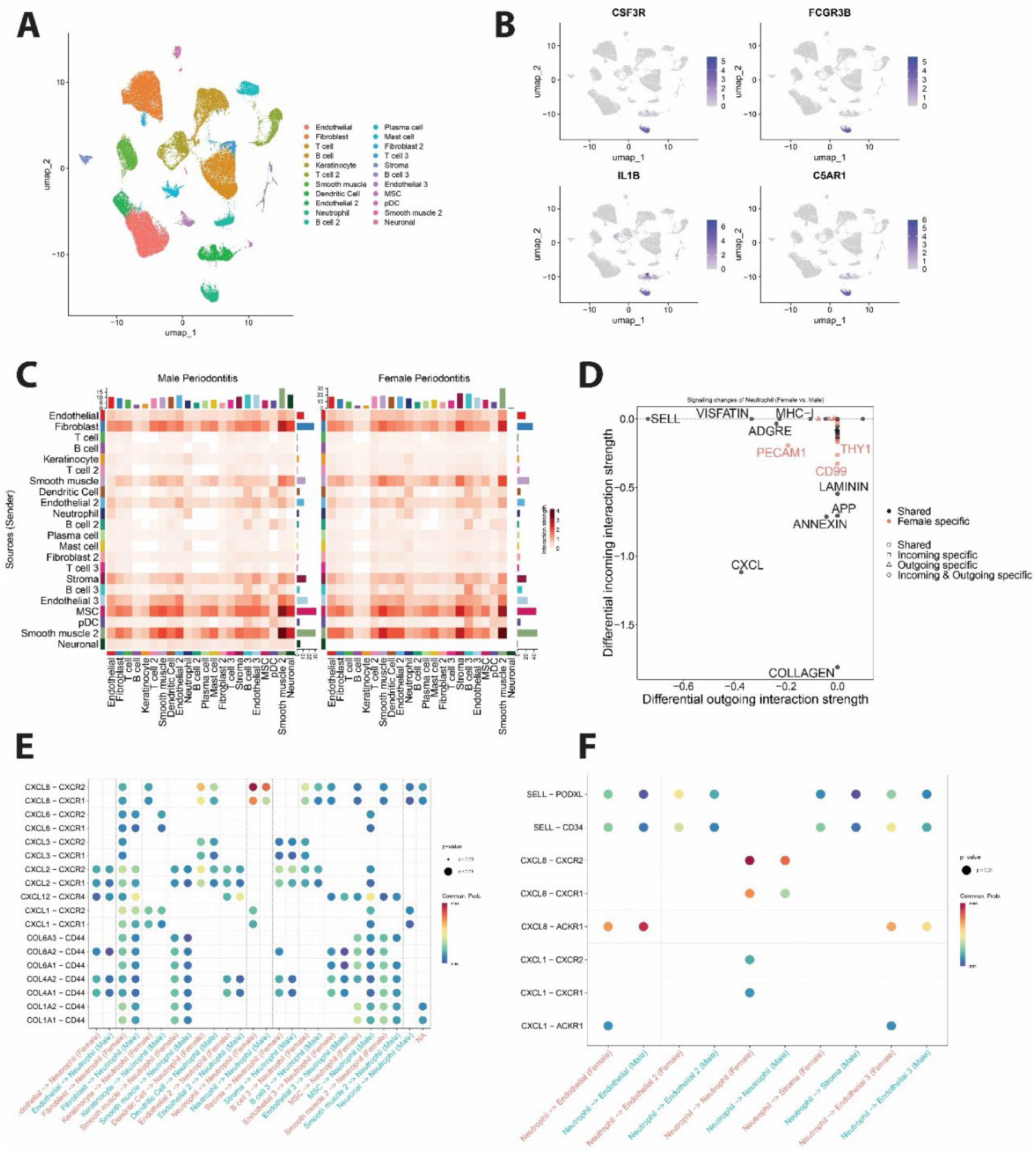
Neutrophil signaling in human periodontitis. A) UMAP plot with labeled clusters from Seurat. B) Heatmap of granulocyte expression markers. C) Signaling information heatmaps from CellChat for male and female patients with periodontitis. D) Differential signaling networks based on incoming and outgoing interaction strength in male and female patients. E) Specific receptor-ligand interactions with neutrophils as the receiver cluster. F) Specific receptor-ligand interactions with neutrophils as the sending cluster.

### Sex differences in neutrophil phenotypes during periodontitis

Reclustering neutrophils identified 10 unique cell clusters (**FIG 5A**). We compared gene expression across the different granulocyte clusters (**FIG 5B, Table S1**). Comparing cluster distributions across conditions found Cluster 0 and Cluster 1 to be significantly associated with periodontitis conditions, with granulocytes from female patients enriched in Cluster 0 and granulocytes from male patients enriched in Cluster 1 (**FIG 5C**). Cluster 0 was enriched for neutrophil markers *S100A9* and *SELL*. Cluster 1 was relatively enriched for *CXCR4* expression with relatively low expression of *SELL*. Gene set enrichment analysis confirmed upregulation of neutrophil-related gene sets in Cluster 0 relative to other granulocyte cells (**FIG 5D**) Cluster 1 was enriched for gene sets related to generalized cytokine signaling in the immune system (**FIG 5E).**

**FIG 5:**
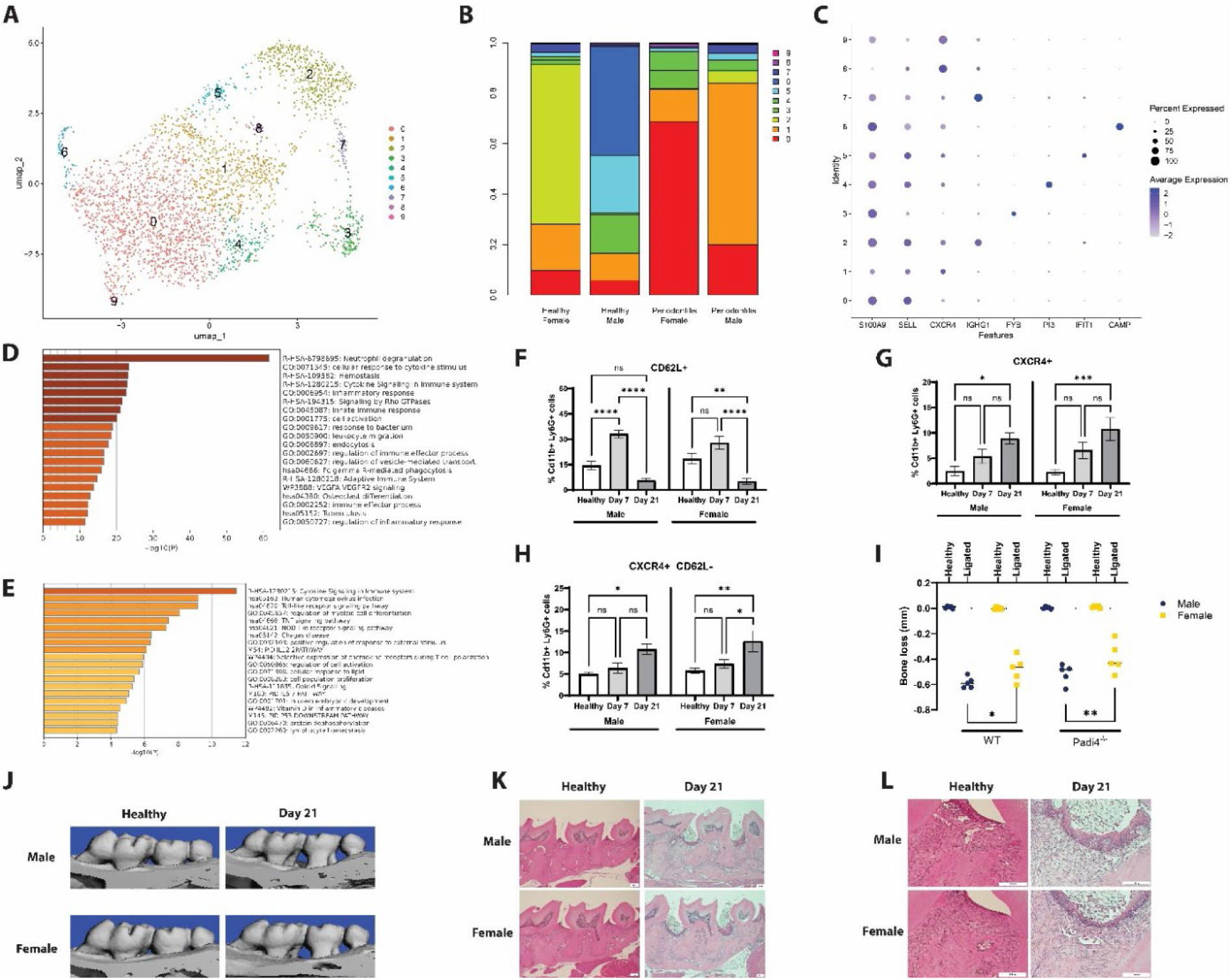
Neutrophil-specific phenotypes in periodontitis. A) Reclustering of neutrophils using Seurat. B) Relative expression of different neutrophil subtypes in healthy and periodontitis male and female patients. C) Marker genes for different neutrophil subtypes in the oral mucosa of males and female with and without periodontitis. D) Gene ontology of upregulated genes from Cluster 0. E) Gene ontology of upregulated genes from Cluster 1. F) Expression of CD62L in neutrophils from the maxillae of mice with different stages of LIP. G) Expression of CXCR4 in neutrophils from the maxillae of mice with different stages of LIP. H) Proportion of CD62L^-^ CXCR4^+^ neutrophils from the maxillae of mice with different stages of LIP. I) Quantification of bone loss after 21 days LIP in Padi4^-\-^ mice J) MicroCT images of maxillae from Padi4^-\-^ mice with LIP. K) Histology of entire ligature site. Scale bar = 100 µM. L) Histology of ligature site. Scale bar = 50 µm. *p<0.05, **p<0.01, ***p<0.001, ***p<0.0001, two-way ANOVA with Tukey’s multiple comparison correction.

Neutrophil extracellular trap (NET) production by infiltrating neutrophils has been previously shown to contribute to bone loss in periodontitis^28^. Since neutrophils contributed significantly to sexual dimorphism in bone loss during periodontitis we investigated the effect of neutrophil phenotype. We examined expression of CD62L (SELL) and CXCR4, two markers of NET-producing neutrophils (**FIG 5F, FIG 5G**). CD62L expression peaked on day 7 in both male and female mice, with no significant difference between males and females (**FIG 5F**). CXCR4 expression peaked at day 21 with no difference between males and females (**FIG 5G**). We additionally characterized “aged” neutrophils, which have been described as CXCR4^+^, CD62L^-^ and are thought to produce more reactive oxygen species and NETs compared with CD62L^hi^ neutrophils^29, 30^. Both male and female mice showed a statistically significant trend towards and increased CD62L^-^ CXCR4^+^ neutrophil fraction over time, with a peak at day 21 (**FIG 5H**). The relative proportion of these cells was the same between male and female mice through the experiment.

We next examined the effects of NET production on sexual dimorphism in periodontitis. This was accomplished by measuring bone loss during periodontitis in Padi4^-\-^ mice, which cannot produce NETs^31^. Male and female Padi4^-\-^ mice both lost statistically significant amounts of bone following ligature placement (**FIG 5I, FIG 5J**), however the amount of bone lost was different between male and female mice. Male Padi4^-\-^ mice had linear bone loss of 0.51 mm compared to 0.39 mm for female Padi4^-\-^ mice (p<0.05). Similar sexual dimorphism was observed between WT and Padi4^-\-^ (**FIG 5I**). Padi4^-\-^ mice did not exhibit as much inflammation in histological characterization compared with WT mice (**FIG 1G, FIG 5K, FIG 5L**). No differences were observed between male and female mice.

## Discussion

Sexual dimorphism of the skeleton is well documented^32^. At maturity, the male skeleton is typically larger and has a higher bone density than the female skeleton. On average, women achieve a lower peak bone mass than men, which is reflected in the clinical finding of 2-4X higher incidence of vertebral fracture and osteoporosis^32^. Mechanisms for these differences, particularly as they relate to alveolar bone, are not completely understood. A variety of factors are responsible for the sexual dimorphism of the skeleton and homeostatic processes of the bone. The organization of bone into functional skeleton is the net result of osteoclasts, which resorb bone, and osteoblasts, which form bone and osteocytes, which coordinate bone development and homeostasis^33^. Differences in skeletal mass or phenotypes between women and men include the action of sex steroids (estrogens and androgens), genetics, environmental/behavior modifiers, and inflammation^34^. Sex hormones, in particular, play a fundamental role in determining both skeletal health as well as bone density-related pathologies that are associated with aging^35–37^. The alveolar bone does not escape the consequences of dimorphic hormone signaling^38^. Sex hormone levels have been associated with the risk of periodontitis in cross-sectional studies^39^, however the biological mechanisms underpinning this risk are not well described. In this study we show differential bone loss resulting from neutrophil-mediated inflammation in male and female mice, suggesting that inherent differences in neutrophils, possibly driven by sex hormone signaling, may provide the basis for this phenotype.

We observed sexual dimorphism in neutrophil-mediated inflammation in the context of periodontal disease in this study. Inflammation is a hallmark of periodontal disease^40^, and discussion of differences between male and female mechanisms of disease progression must include innate immune cells. Toll-like receptor 7 (TLR7) is encoded on the X chromosome and may escape X-inactivation in certain cell types, leading to higher levels of the TLR7 in female cells relative to male counterparts^41, 42^. Toward this end, TLR signaling pathways yield sexual dimorphic responses including higher cytokine production in females of myeloid primary response gene 88 (MYD88), retinoic acid-inducible gene-I (RIGI), interferons, Janus kinase 2(JAK2), signal transducer and activator of transcription 3 (STAT3), NF-κB, interferon gamma (IFN-γ), and tumor necrosis factor alpha (TNF)^43^. Compared to females, peritoneal macrophages from males express higher levels of TLR4, which is a receptor for bacterial wall lipopolysaccharides (LPS) that induces large-scale cellular inflammatory responses in the oral mucosa with more robust levels of CXCL10^44^. Notably, female macrophages have enhanced phagocytosis and antigen presentation capacity to T-lymphocytes for the initiation of the adaptive immune response^44^. Sexual dimorphism during immune activation has been well established, however reports defining the mechanistic role these differences play in the progression of periodontal disease are limited. A model of periodontitis induced by *A. actinomycetemcomitans*-derived LPS enhanced RANKL-induced osteoclastogenesis in male mice relative to female mice^45, 46^. The genes *Nfatc1* and *Tm7sf4* (encoding dendritic cell-specific transmembrane protein or DCSTAMP) were also more highly expressed in male osteoclasts in this model, suggesting an innate immune driven mechanism for these differences^45, 46^. In our studies, innate immune activation appeared to play a role in sex-specific differences in hematopoiesis during chronic periodontitis.

The periodontal structures provides a unique platform for the mutualistic relationship between the host and indigenous microbial community, which maintains immunological tone in homeostatic conditions^47^. Neutrophils represent the most abundant innate immune cell at oral mucosal tissues and barrier sites such as the gingiva^22^. Neutrophils are key mediators of health and disease in the periodontal space within the oral cavity. In their absence or when they are functionally defective, as is the case in certain congenital disorders, severe forms of periodontitis develop at early ages^48^. However, the presence of supernumerary or hyper-responsive neutrophils, leads to imbalanced host-microbe interactions that culminate in dysbiosis and inflammatory tissue breakdown^49^. In this study, depletion of neutrophils resulted in a reversal of the sexual dimorphic bone loss phenotype observed in healthy mice, with female mice losing more bone than male mice (**FIG 3**). This result possibly indicates differences in neutrophil function and susceptibility to diseases related to neutrophil deficiency in males and females, however this relationship remains undefined.

There are sex-biased differences in innate, humoral, and cellular immune recruitment and function in humans; however, the exact mechanisms that drive this notable contrast remain insufficiently characterized^50^. Young, but not older, healthy adult males displayed significant enhancement in immature neutrophil gene signatures compared to age-matched females, resulting in 1) less activated male neutrophil phenotype based on cell surface markers, 2) hampered response to cytokine stimulation, and 3) decreased ability to form neutrophil extracellular traps (NETs) following stimulus^51^. Conversely, healthy young adult females have an activated/mature neutrophil profile characterized by enhanced type I IFN pathway activity and enhanced proinflammatory responses^52^. We specifically examined one pathway, production of neutrophil extracellular traps, on sexual dimorphism in periodontal disease. We observed that this function of neutrophils did not significantly impact differences in bone loss between male and female mice, indicating other functions such as production of reactive oxygen species of other proinflammatory cytokines are likely driving this phenotype.

## Conclusion

Sexual dimorphism can be observed in the clinical phenotype of experimental periodontitis. Differences in bone loss are driven, in part, by differences in neutrophil recruitment and phenotype at the site of inflammation. Future therapies could target these differences towards the goal of personalizing treatment for periodontal disease.

## Acknowledgements

This work was funded by NIH grant R00DE029756 (AMD) and research funds from the University of Michigan School of Dentistry (JTD). VM is funded by NIH grant K12DE027826.

---

**FIG S1:**
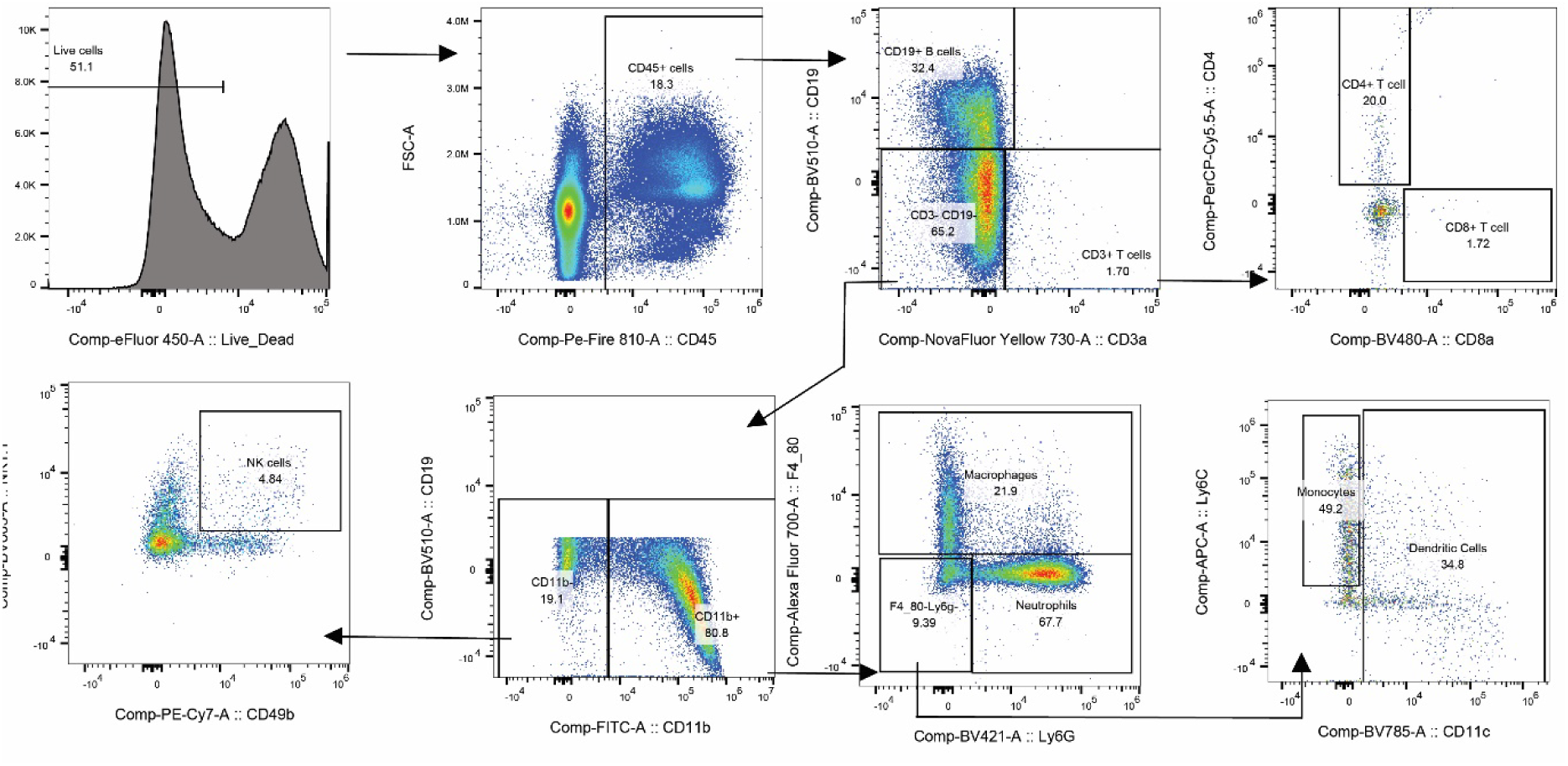
Flow cytometry gating to identify immune cell populations.

